# Spiral Waves and Turbulence in Mathematical Models of Oncolytic Virotherapy

**DOI:** 10.64898/2026.01.01.697297

**Authors:** Tejas Bansod, Thomas Hillen

## Abstract

Oncolytic virotherapy is a promising targeted cancer treatment that employs viruses, which selectively infect tumor cells. Although its clinical efficiency has remained limited and it is often used in conjunction with other therapies, advances in genetic engineering have produced stronger and more selective viral strains, prompting continued interest in their dynamics. In particular, previous studies have noted that viruses with sufficiently high replication rates can induce oscillations reminiscent of predator–prey systems. Here, we extend this analysis to the spatial domain by starting from an established tumor–virus reaction–diffusion model, performing a center-manifold reduction that incorporates nonlinear terms to derive a complex Ginzburg–Landau amplitude equation, and estimating its parameters directly from the original kinetics. This reduced normal form equation explains the emergence of experimentally observed patterns — such as hollow rings and target waves — and shows that, at longer timescales, these patterns naturally evolve toward spiral waves and a turbulent regime. Our work provides a mechanistic link between the kinetic Hopf bifurcation and the rich spatiotemporal structures observed in oncolytic virotherapy models, suggesting that these patterns are not numerical artifacts but an intrinsic feature of the system.

## 1 INTRODUCTION

Cancer, defined by the uncontrolled growth and spread of abnormal cells, remains one of the leading causes of death worldwide [32]. Over the years, diverse treatments have been devised —chemotherapy [11], radiotherapy [16], immunotherapy [23], and more recently, oncolytic virotherapy [28]. Oncolytic virotherapy employs replicating viruses, which selectively infect and destroy tumor cells while sparing healthy tissues [22]. It looks very appealing in principle but in practice there are countless uncertainties at the biological level [35]. For example, how does the immune system react to this foreign virus[30]? How easily can tumor develop viral resistance [6]? These type of questions arise for any treatment in the oncologist’s arsenal, but fortunately these uncertainties can be made more tangible by formulating them as mathematical questions, providing at least a benchmark for expected outcomes and make an educated guess about directions, which promise the most improvement [3].

If we adopt an ecological picture, we have oncolytic virus as a parasite trying to invade the ecosystem of the tumor host [29]. Host-parasite systems have been studied extensively, particularly in the models of epidemiological outbreaks [14] which highlight the basic reproduction number (*R*_0_) as the main parameter to measure the contagiousness and the effect of the virus [10]. For oncolytic virotherapy, a similar approach has been taken to formulate Ordinary Differential Equations (ODEs) [5, 34] which identify a replication threshold beyond which the tumor population is expected to show oscillations. In this paper, we start from a previous model [3], which demonstrated that for high enough viral replication, these viral infection oscillations can drive the tumor to near-extinction. We note however that current viruses are still far from this Hopf bifurcation threshold [3]. Therefore, the natural question to ask here is: even if virus replication could be increased as much as desired, would that be enough for effective tumor eradication [27]?

An important aspect of the viral infection of cancer is the spatial extent of the tumor. Tumors have a finite size and the virus cannot reach all cells instantly, it needs time to infect, produce more virus, infect again etc. As a result periodic dynamics at different location along the tumor mix and exchange, leading to interest ting spatio-temporal patterns. [3]. Moreover, both in vitro and in silico approaches [40, 6] have demonstrated that the spatial structure of the tumor is crucial for therapeutic success. In particular, Wodarz et al. [40] used spatial models to show that the dynamics driven by waves of virus chasing the tumor lead to three distinct outcomes — hollow rings (successful clearance), concentric rings (tumor regrowth), and fragmented aggregates.

The previous model by Baabdulla et. al [3] consisted of three components - uninfected cancer, infected cancer and free virus (C,I and V respectively) where virus diffuses more than the tumor due to its smaller size [3]. By simulating this Reaction-Diffusion system, the patterns observed before by Wodarz. et al [40] could be reproduced. In this work, we aim to demonstrate that these patterns are robust and not merely artifacts of some specific parameter choices. Our approach is to use mathematical reductions to make clear the basic mechanism underlying these patterns. We present a flow chart for our reduction steps in Figure 1.

**Figure 1.**
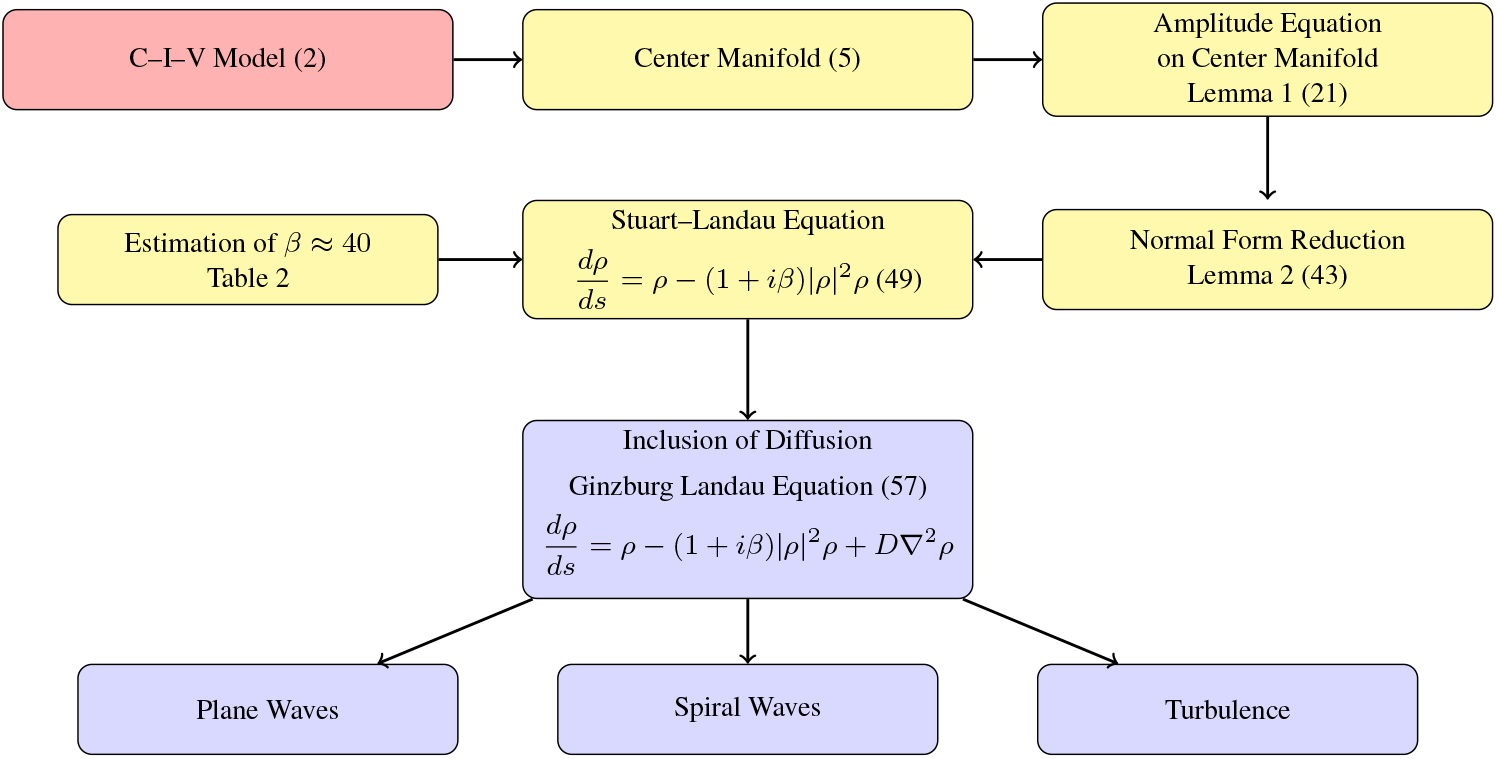
Flow of analytical reductions from center–manifold dynamics to amplitude equations, parameter estimation, and pattern formation. The steps related to the well mixed ODE system are highlighted in yellow, while the results for the spatial PDE are shown with blue background

In this approach, we first reduce the temporal dynamics of the full model to its center manifold [8] just above the Hopf bifurcation threshold. This reduction leads to a two-dimensional system, which can be related back to the original model and whose linear dynamics are directly given by the complex pair of eigenvalues. Further, to capture the resultant limit cycle in this reduced system, we determine the necessary nonlinear terms for the center manifold dynamics, which turn out to be quadratic since the original model’s nonlinearities are also purely quadratic. However, since these nonlinear terms are coordinate dependent, we perform a normal form transformation through a lemma by Kuznetzsov [20] which allows us to interpret the reduced system in terms of the Stuart–Landau Equation (SLE) [21]. This reduction constitutes the first analytical part of the paper, reiterating the well-known result that systems near a supercritical Hopf bifurcation can be universally represented by the SLE via normal form theory [17]. Upon non-dimensionalization, this results in a single asynchrony parameter *β* [19], which is a measure of frequency shifts between oscillations of varying amplitudes, and which we estimate numerically for our system.

After completely understanding the kinetic behavior in its normal form (the SLE), we proceed to the second part of this work, where the role of diffusion is concerned. This spatiotemporal extension gives the well-known Complex Ginzburg–Landau Equation (CGLE) [2] as normal form. The CGLE is known to generate a spectacular range of dynamic patterns, including plane waves, spiral waves, and spatiotemporal turbulence [19]. Spiral waves, for instance, are among the most striking patterns observed across diverse systems, from the Belousov–Zhabotinsky reaction [42] to electrical activity in cardiac tissue [38]. However, our numerical estimate of the parameter *β* suggests that the C-I-V system operates well beyond the regime that can support stable spiral waves. Consequently, turbulent patterns appear to be the norm rather than the exception for a large range of biological parameters in our system.

This work proceeds as follows: Section 2 introduces the C-I-V model and recalls the bifurcation structure as derived in the literature [3, 1, 27]. In Section 3 we focus on the ODEs of the well mixed version (no diffusion) and reduce the system near the Hopf bifurcation on the 2-dimensional center manifold to a complex Stuart-Landau equation. In Section 4 we add back diffusion and explain how the diffusion terms are carried through the reduction steps, leading to the complex Ginzburg-Landau equation (CGLE) as normal form. We show some numerical simulations for the full (*C, I, V*) model, followed by some simulations for spiral waves and their transition to turbulence for the CGLE. Finally, in Section 5, we conclude by discussing the biological implications of the turbulent behavior, the role for other factors such as immune system and resistance, and offer a heuristic explanation for periodic spot splitting phenomenon observed in an earlier work [3].

## 2 MODEL DESCRIPTION AND OSCILLATIONS VIA HOPF BIFURCATION

Oncolytic viruses are engineered to exploit the differential susceptibility of tumor and normal cells. To capture the essential feedback between viral replication and tumor regrowth, we employ the nondimensionalized C–I–V model of Baabdulla and Hillen [3], which extends the framework of Pooladvand et al. [27]. The model describes three interacting densities: *C*(*t*, **x**): uninfected tumor cells, *I*(*t*, **x**): infected tumor cells and *V* (*t*, **x**): free virus particles. Their dynamics follow a system of reaction–diffusion equations:

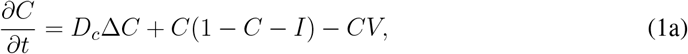

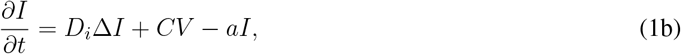

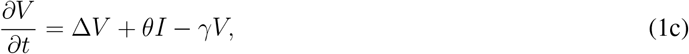

where Δ denotes the Laplace operator, and *D*_*c*_, *D*_*i*_ are the diffusion coefficients for cancer cells and infected cancer cells, respectively. Tumor proliferation is described through a logistic term *C*(1 − *C* − *I*) limited by space and resources. Viral infection is described by a mass-action term − *CV*, infected cells lyse at rate *a*, and produce new virions with rate *θ*. Viral particles are cleared with rate *γ*. Most parameters are normalized to equal 1, except the death rate of infected cells *a*, the removal rate of free virus particles *γ*, and the burst size *θ*. The dimensional version of the model, together with realistic parameter estimates and sensitivity analysis can be found in the cited literature [3, 27]. Together, these terms form a minimal yet complete model of virus infection-mediated tumor control, balancing growth, lysis, and diffusion.

It should be noted that we exclude two more terms, which are often included in these oncolytic virus models, see for example Al-Tuwairqi *et al*. [1]. We ignore a viral loss term of the form − *β*_2_*CV* in the last equation, since we assume, as in [39, 13, 41, 3], that the loss of virus particles due to an infection of a cell is minimal. Moreover, we do not include a natural death term for cancer cells. A death term in the *C* equation will shift the value of the Hopf bifurcation point, but it will not change the dynamics near that Hopf point.

We study (1) on a two–dimensional domain Ω ⊂ ℝ^2^ with zero–flux boundaries,

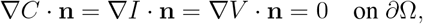

so that both cells and virions remain confined within Ω. This closed-domain setting mirrors in-vitro tumor spheroid experiments and petri-dish experiments [4], where viral spread occurs within a constrained microenvironment.

We follow [3] in adopting the values listed in Table 1. Among these, the viral production rate *θ* acts as the primary bifurcation parameter controlling infection strength and oscillatory onset.

**Table 1.** Non-dimensional parameter set used in [3].

| Parameter | Description | Value |
| --- | --- | --- |
| $D_c$ | diffusion of uninfected cells | 0.025 |
| $D_i$ | diffusion of infected cells | 0.025 |
| $a$ | death rate of infected cells | 3.33 |
| $\gamma$ | viral clearance rate | 13.33 |
| $\theta$ | effective viral production rate | [0, 500] |

### 2.1 Kinetic Behavior

To understand the mechanisms driving oscillations, we first suppress spatial dependence and analyze the corresponding well-mixed system:

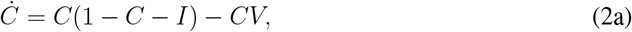

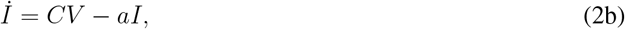

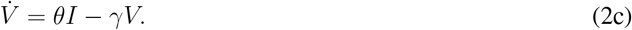

This reduction highlights the feedback loop between viral amplification and tumor recovery. System (2) has been analysed previously, see for example in [27, 3] and we summarize the main properties here. The system admits three equilibria:

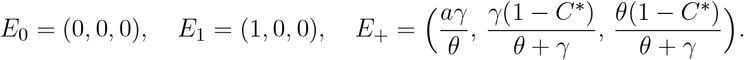

The coexistence equilibrium *E*_+_ exists for *θ* > *aγ* and is born through a transcritical bifurcation at *θ*_*T*_ = *aγ*. Biologically, this threshold marks the point where viral replication compensates for clearance, allowing infection to persist. As *θ* increases, *E*_+_ eventually loses stability through a Hopf bifurcation at *θ*_*H*_, producing sustained oscillations in tumor and viral populations. We show the bifurcation diagram in Figure 2, where *θ* is the bifurcation parameter and all other parameters are fixed as in Table 1. For three values of *θ* we show the time dynamics of *C*(*t*) in Figure 3.

**Figure 2.**
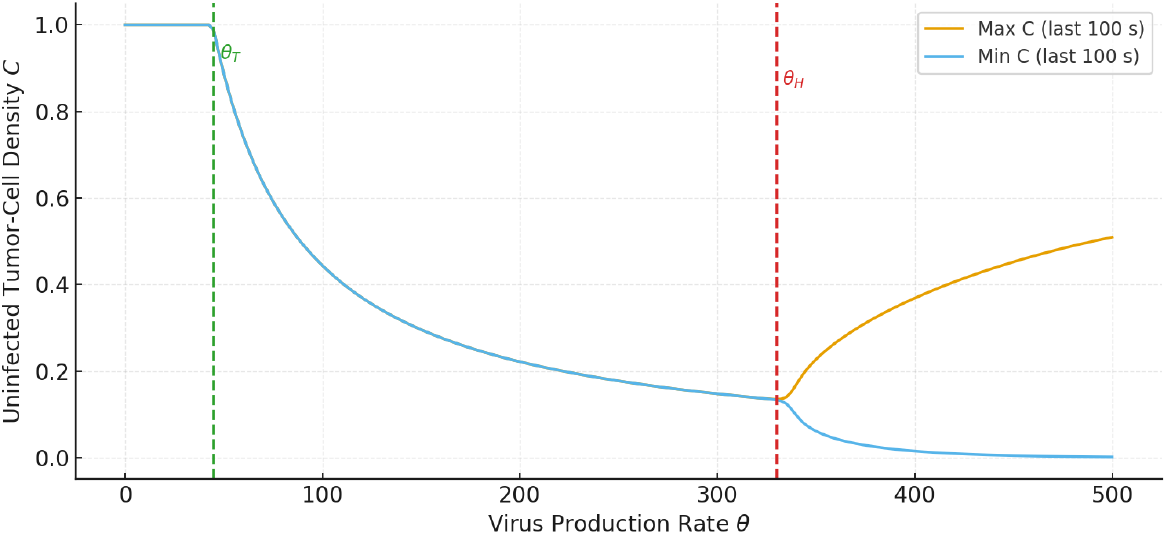
Kinetic bifurcation diagram versus *θ* for (2): creation of the coexistence state at the transcritical threshold *θ*_*T*_ and loss of equilibrium stability at the Hopf threshold *θ*_*H*_

**Figure 3.**
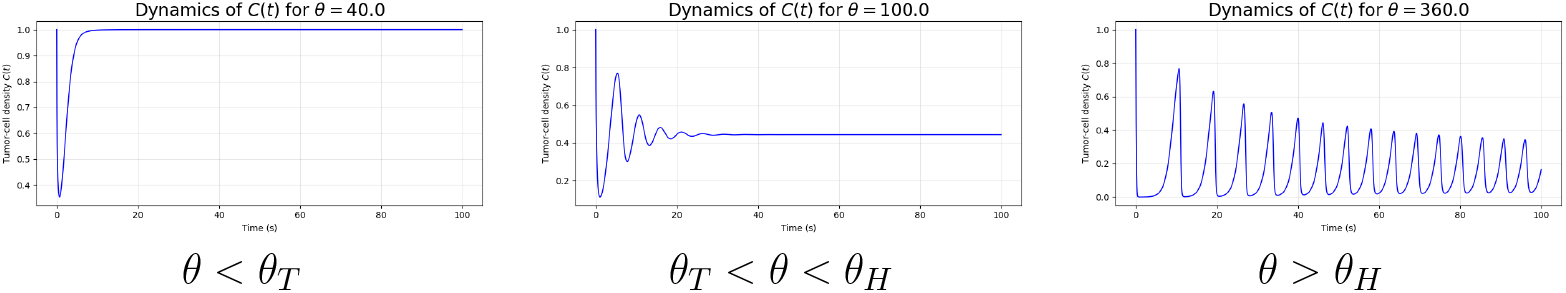
Temporal dynamics of *C*(*t*) across parameter regimes of (2). (a) tumor persistence; (b) stable coexistence; (c) sustained oscillations following the Hopf bifurcation.

The oscillations observed in Figure 3 originate from a loss of stability of the coexistence equilibrium *E*_+_. To determine the mechanism, we linearize the kinetic system (2) about *E*_+_ and examine the eigenvalues of its Jacobian matrix:

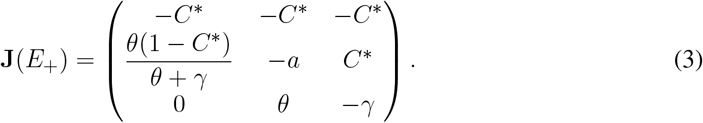

The characteristic polynomial of **J**(*E*_+_) depends smoothly on the viral production rate *θ*. At a critical value *θ* = *θ*_*H*_, one pair of complex conjugate eigenvalues *λ*_2,3_ = ± *iω*_0*H*_ crosses the imaginary axis, while the remaining real eigenvalue *λ*_1_ stays strictly negative. Baabdulla and Hillen [3] proved that this crossing satisfies the standard conditions of a forward Hopf bifurcation.

Moreover, near *θ*_*H*_, the system’s behavior is governed by the two-dimensional eigenspace associated with the critical pair *λ*_2,3_, while the remaining direction is rapidly damped (*λ*_1_ < 0.). This separation of timescales allows reduction to a planar oscillatory subsystem on the corresponding center manifold.

## 3 ANALYSIS AND MODEL REDUCTIONS

We perform a systematic model reduction to obtain a Stuart-Landau normal form. We proceed as follows:

### 3.1 Reduction to the Centre Manifold

At the Hopf threshold, the Jacobian **J**(*E*_+_(*θ*_*H*_)) possesses a purely imaginary eigenpair ±*iω*_0_ and a single stable eigenvalue *λ*_1_(*θ*_*H*_) *<* 0. Let **e**_3_ denote the stable eigenvector and 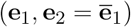 the conjugate pair spanning the oscillatory subspace. The dynamics decompose accordingly: fast decay along **e**_3_ and sustained oscillations within span {**e**_1_, **e**_2_ }. To make this decomposition explicit, we first shift the equilibrium to the origin:

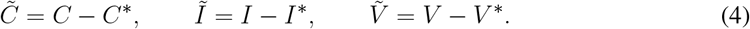

Perturbations 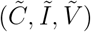 are then expressed in the eigenspace basis as span {**e**_1_, **e**_2_} to leading order as

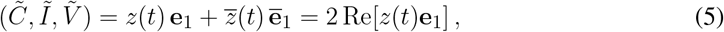

introducing the complex amplitude *z*(*t*) = *u*(*t*) + *iv*(*t*). The real variables (*u, v*) represent the reduced coordinates on the center manifold and form the foundation of the Stuart–Landau reduction carried out later.

We make this transformation explicit, by computing the eigenvectors **e**_1_ and **e**_2_. We normalize the eigenvector **e**_1_ so that its third component is real and equal to 1. Then it has the general form

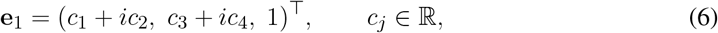

where the coefficients *c*_1_, *c*_2_, *c*_3_, *c*_4_ will be found later. Substituting (6) into (5) gives

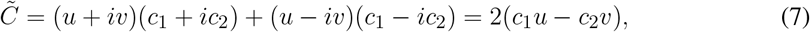

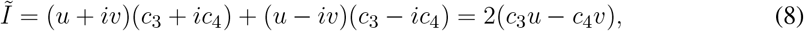

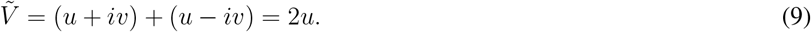

which defines the forward mapping 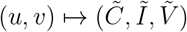 with

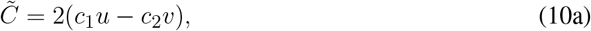

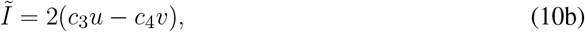

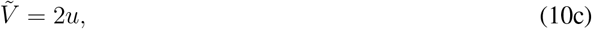

Solving for (*u, v*) yields

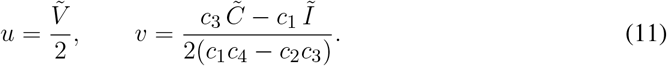

Biologically, *u* corresponds to the viral component, while *v* represents a weighted average of the tumor components.

To determine **e**_1_ explicitly, we solve

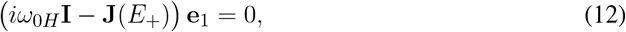

with **J**(*E*_+_) given in (3). Setting **e**_1_ = (*l*_1_, *l*_2_, 1)^⊤^ gives the linear system

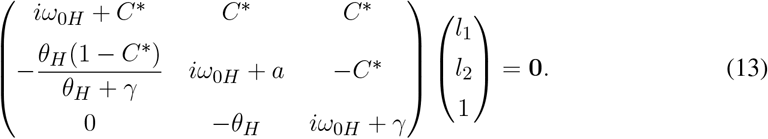

From the third row we obtain

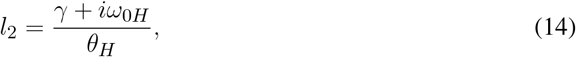

and substituting into the first row gives

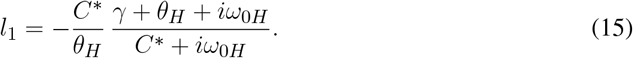

Thus,

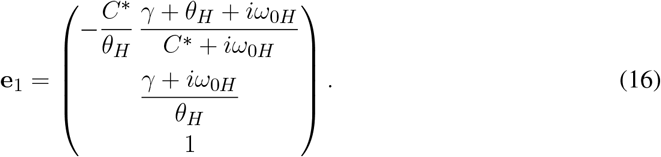

Writing *l*_1_ = *c*_1_ + *ic*_2_ and *l*_2_ = *c*_3_ + *ic*_4_ yields the real coefficients

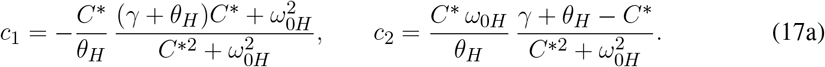

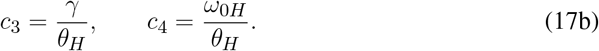

Substituting into (11) provides the full coordinate transformation between the physical variables (*C, I, V*) and the reduced oscillatory variables (*u, v*). In particular, the denominator in the expression for *v* is

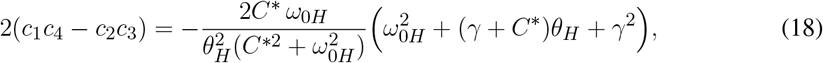

which is nonzero at the Hopf point, confirming that the transformation is well defined in a neighborhood around the bifurcation point.

### 3.2 Nonlinear Dynamics on the Centre Manifold

For *θ* near the Hopf threshold *θ*_*H*_ we write the eigenvalue associated with **e**_1_ in the general form

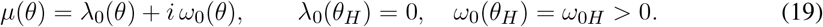

The real part *λ*_0_(*θ*) governs the growth or decay of oscillations, while *ω*_0_(*θ*) sets their frequency. On the center manifold, using the complex coordinate *z* = *u* + *iv* from (5), the leading-order linear dynamics take the form

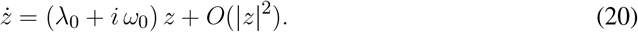

We show that the nonlinear terms are quadratic.

#### Lemma 1.

*The dynamics on the centre manifold at the Hopf point are given by the nonlinear complex amplitude equation*

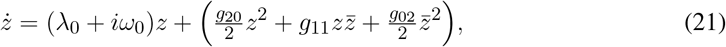

*with coefficients as defined below in(25,26,30)*.

#### Proof.

The original C–I–V model (2) contains only quadratic nonlinearities (terms such as *C*^2^, *CI, CV*). After translation to *E*_+_, the constant terms vanish, leaving a purely quadratic remainder. Since the change of variables to (*u, v*) in (11) is linear, this degree is preserved.

Hence, the reduced vector field on the center manifold is quadratic:

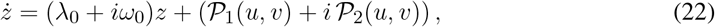

with

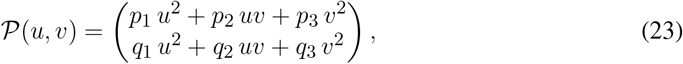

for some coefficients *p*_*i*_, *q*_*i*_ ∈ ℝ.

Expanding *u*^2^, *v*^2^, and *uv* in powers of *z* = *u* + *iv* and 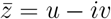 gives

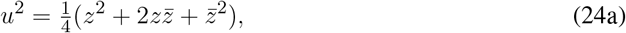

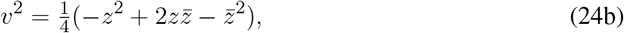

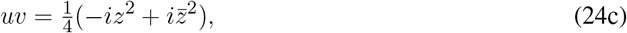

which leads to the compact complex form (21). with coefficients

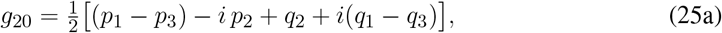

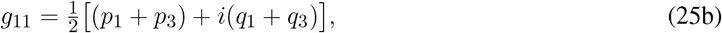

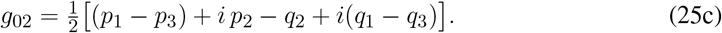

It remains to determine the coefficients *p*_*i*_ and *q*_*i*_ at the Hopf point. By construction, 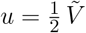, and from (2) the 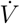-equation is purely linear,

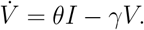

Using 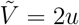 and *Ĩ* = 2(*c*_3_*u* − *c*_4_*v*) from (10) gives

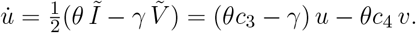

Substituting *c*_3_ = *γ/θ* and *c*_4_ = *ω*_0*H*_*/θ* Yields 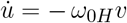, confirming that

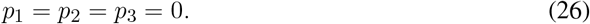

To find the coefficients *q*_1_, *q*_2_, *q*_3_, we consider the parametrization of **e**_1_ = (*c*_1_ + *ic*_2_, *c*_3_ + *ic*_4_, 1)^⊤^. Recall from (10) that

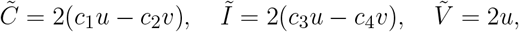

and from (11),

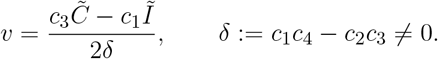

The quadratic parts of the ODE system (2) in shifted variables are

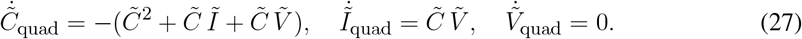

Because *c*_1_, *c*_3_, *δ* are constants,

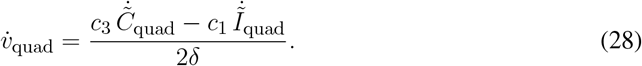

Expressing 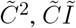, and 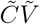 in terms of (*u, v*) and simplifying yields

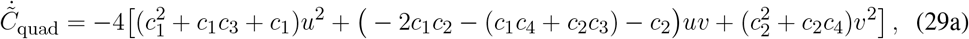

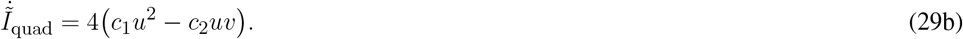

Substituting (29) into (28) and matching terms with 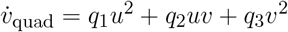 yields

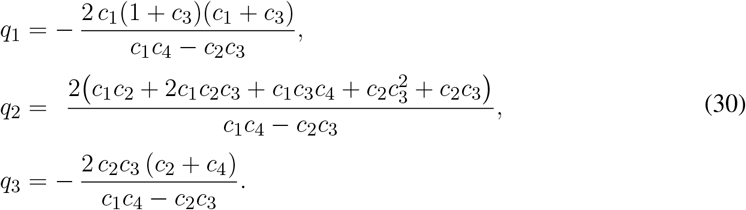

#### Remark 1.

*At θ* = *θ*_*H*_, *these coefficients fully characterize the quadratic nonlinearity of the reduced oscillatory system. While their continuation beyond θ*_*H*_ *is nontrivial, we will see later that explicit values are unnecessary for deriving the normal form and assessing pattern formation in the spatially extended model*.

### 3.3 Normal Form Reduction

To pass from the quadratic center–manifold system (21) to a universal amplitude description, we remove quadratic terms in (21) by a parameter-dependent near-identity change of variables, following the Hopf normal-form construction (cf. Kuznetsov [20], Lemma 3.4) with bifurcation parameter *θ*. (Throughout, the coefficients *p*_*i*_(*θ*), *q*_*i*_(*θ*) and hence *g*_*ij*_(*θ*) depend smoothly on *θ*.)

#### Lemma 2

(Removal of quadratic terms near Hopf). *Consider the complex amplitude equation on the center manifold*

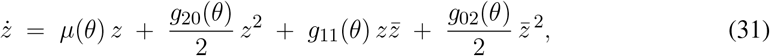

*where µ*(*θ*) = *λ*_0_(*θ*) + *iω*_0_(*θ*) *with λ*_0_(*θ*_*H*_) = 0 *and ω*_0_(*θ*_*H*_) = *ω*_0*H*_ > 0. *There exists an invertible, θ-dependent near-identity change of complex coordinate*

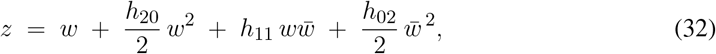

*valid for sufficiently small* |*w*| *and* |*θ* − *θ*_*H*_ |, *which transforms* (31) *into an equation without quadratic terms:*

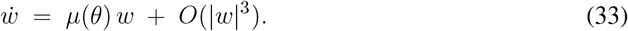

#### Proof.

The idea is to remove quadratic terms in (31) by a near-identity change of variables (32), where the constants *h*_20_, *h*_11_, *h*_02_ will be determined at a later stage of the proof.

We begin from the near-identity change of variables defined in (32), which upon inverting formally gives

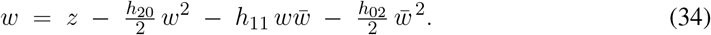

Since *z* = *w* + *O*( |*w*|^2^), we may replace each quadratic monomial in *w* by its *z*–counterpart up to cubic error:

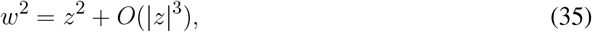

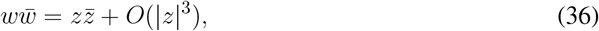

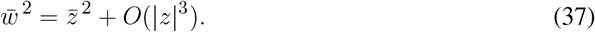

Substituting (35)–(37) into (34) yields

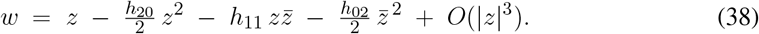

Next, we want to use (31) to write dynamical equation for 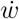: We start by differentiating (38):

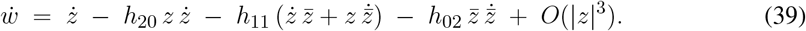

And next we substitute 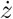 from (31), keeping only terms up to quadratic order. Since

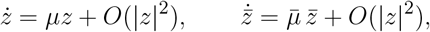

(39) becomes

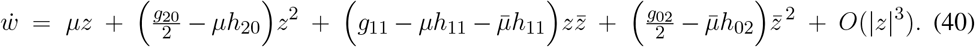

Finally, using *z* = *w* + *O*(|*w*|^2^) and collecting quadratic terms in *w*, we obtain

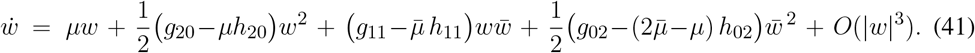

Thus, by choosing

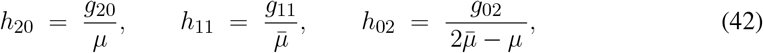

we cancel all quadratic terms and obtain the cubic normal form (33). At *θ* = *θ*_*H*_ we have *μ* (*θ*_*H*_) = *iω*_0*H*_ ≠ 0 and 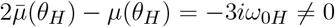, so the denominators remain nonzero for |*θ* − *θ*_*H*_| sufficiently small.

Consequently, applying Lemma 2 to (21) gives the cubic **Stuart–Landau normal form**

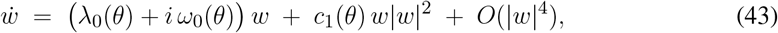

where *c*_1_(*θ*) is to be determined by the original C–I–V parameters.

### 3.4 Phase and Amplitude Equations

We now reinterpret the Stuart–Landau normal form

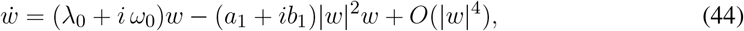

where *λ*_0_(*θ*_*H*_) = 0 and *ω*_0_(*θ*_*H*_) = *ω*_0*H*_ > 0, and where *c*_1_(*θ*) = − *a*_1_ − *ib*_1_ is the cubic normal form coefficient.

Writing the complex amplitude as *w* = *re*^*iϕ*^ with *r* ≥ 0 and separating real and imaginary parts yields

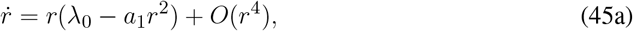

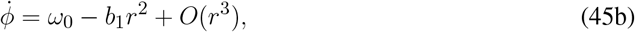

which, to leading order, are the classical Λ−Ω *equations* [18, 19] with

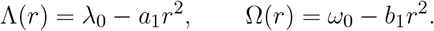

For a supercritical Hopf bifurcation (*a*_1_ > 0), the amplitude equation admits a stable limit cycle of radius 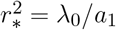, oscillating at frequency 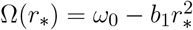.

To remove the trivial rotation at frequency *ω*_0_, we introduce the co-rotating variable

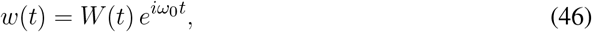

so that |*w*| = |*W*|. Differentiation and substitution into (44) give

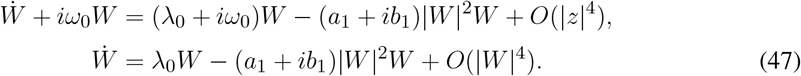

Assuming *a*_1_ > 0, we scale variables to normalize the linear and cubic terms by defining

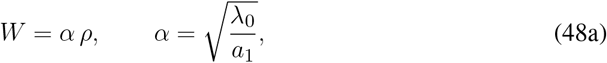

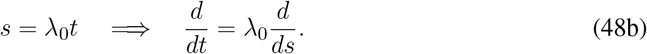

Using *α*^2^ = λ_0_*/a*_1_ and substituting into (47) yields the normal form

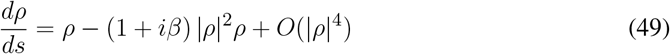

where

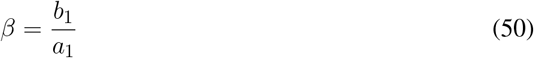

captures the strength of amplitude–frequency coupling.

In this nondimensional form, the amplitude saturates at |*ρ*| = 1 while the imaginary coefficient *β* |*ρ*|^2^ generates the amplitude-dependent frequency shift characteristic of oscillatory media near Hopf bifurcation.

### 3.5 Evaluating the Normal –Form Parameter *β*

The parameter *β* in (49) quantifies the nonlinear frequency shift per unit amplitude growth and links the reduced dynamics to the underlying C–I–V kinetics. To express it in measurable terms, note that for *θ* > *θ*_*H*_ the stable limit cycle with radius *r*_∗_ satisfies

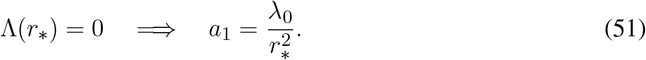

Its oscillation frequency,

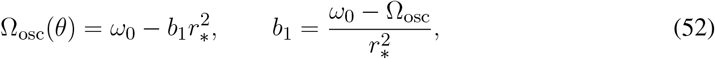

then gives

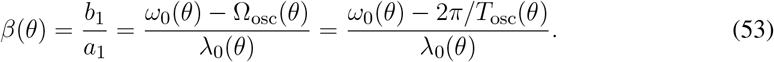

Thus *β* can be obtained directly by combining the linear eigenvalues (*λ*_0_, *ω*_0_) of the coexistence state with the nonlinear oscillation period *T*_osc_(*θ*) of the full system.

This observation leads to the following workflow: For each *θ* near the Hopf threshold:

1. Compute the leading complex eigenvalue *µ*(*θ*) = *λ*_0_(*θ*) + *iω*_0_(*θ*) of the Jacobian (3).

2.Integrate the ODE (2) until transients decay and measure the oscillation period *T*_osc_(*θ*) from successive peaks of *C*(*t*).

3.Evaluate *β*(*θ*) via (53).

**Example:** For our base parameter set from Table 1 we show in Figure 4 the eigenvalue pair of *E*_+_ as *θ* increases: *λ*_0_(*θ*) crosses zero at *θ*_*H*_, while *ω*_0_(*θ*) varies slowly. Figure 5 illustrates the resulting oscillations of *C*(*t*) for representative values of *θ* above the threshold. Combining linear and nonlinear data gives the nearly constant *β*(*θ*) values in Table 2. The ratio *ω*_0_ − Ω_osc_ grows roughly proportionally to *λ*_0_, keeping *β* of order 40 with only weak dependence on *θ*. This suggests that the frequency–amplitude coupling is an intrinsic property of the underlying C–I–V kinetics rather than a parameter-specific artifact.

**Table 2.** Comparison of Jacobian eigenvalues and measured oscillation properties for *θ* = 340–360. The synchrony parameter *β*(*θ*) is computed from (53). Note the near constancy of *β* across the range.

| $\theta$ | $\lambda_0$ | $\omega_0$ | $T_{\text{osc}}$ | $\Omega_{\text{osc}}$ | $\beta$ |
| --- | --- | --- | --- | --- | --- |
| 340 | 0.0004 | 1.5161 | 4.191 | 1.499 | 43.0 |
| 341 | 0.0006 | 1.5164 | 4.217 | 1.490 | 44.0 |
| 342 | 0.0009 | 1.5167 | 4.235 | 1.484 | 37.6 |
| 343 | 0.0011 | 1.5170 | 4.259 | 1.475 | 38.2 |
| 344 | 0.0013 | 1.5174 | 4.286 | 1.466 | 39.1 |
| 345 | 0.0015 | 1.5177 | 4.309 | 1.458 | 39.5 |
| 346 | 0.0018 | 1.5180 | 4.332 | 1.450 | 38.0 |
| 347 | 0.0020 | 1.5183 | 4.359 | 1.441 | 38.6 |
| 348 | 0.0022 | 1.5186 | 4.382 | 1.434 | 39.3 |
| 349 | 0.0024 | 1.5190 | 4.409 | 1.425 | 39.2 |
| 350 | 0.0026 | 1.5193 | 4.432 | 1.418 | 39.0 |
| 351 | 0.0029 | 1.5196 | 4.455 | 1.410 | 38.3 |
| 352 | 0.0031 | 1.5199 | 4.481 | 1.402 | 38.1 |
| 353 | 0.0033 | 1.5202 | 4.505 | 1.395 | 37.7 |
| 354 | 0.0035 | 1.5205 | 4.529 | 1.387 | 37.5 |
| 355 | 0.0037 | 1.5208 | 4.557 | 1.379 | 37.0 |
| 356 | 0.0039 | 1.5211 | 4.580 | 1.372 | 36.9 |
| 357 | 0.0041 | 1.5214 | 4.605 | 1.364 | 36.6 |
| 358 | 0.0043 | 1.5217 | 4.629 | 1.357 | 36.4 |
| 359 | 0.0045 | 1.5220 | 4.655 | 1.350 | 36.0 |
| 360 | 0.0047 | 1.5223 | 4.680 | 1.343 | 36.2 |

**Figure 4.**
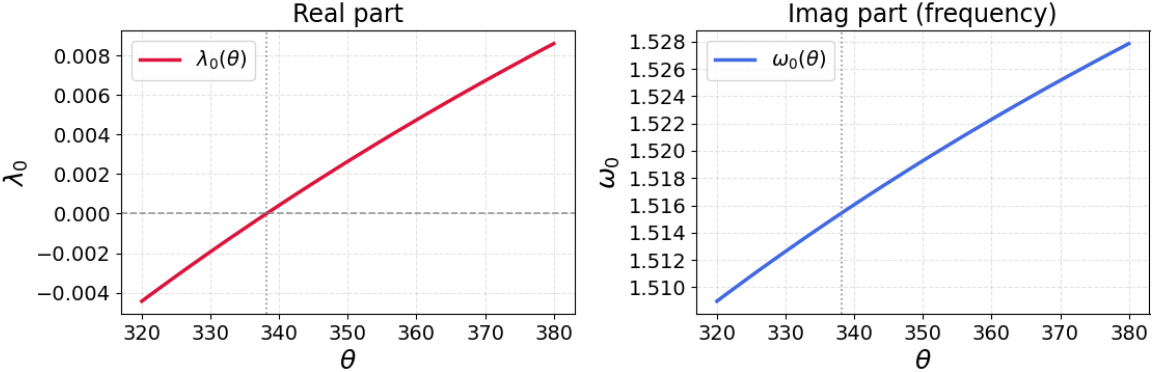
Leading eigenvalue components of the coexistence equilibrium *E*_+_ (**??**) as functions of viral production rate *θ*. The real part *λ*_0_(*θ*) crosses zero at *θ* = *θ*_*H*_, marking the Hopf bifurcation, while the imaginary part *ω*_0_(*θ*) varies only weakly.

**Figure 5.**
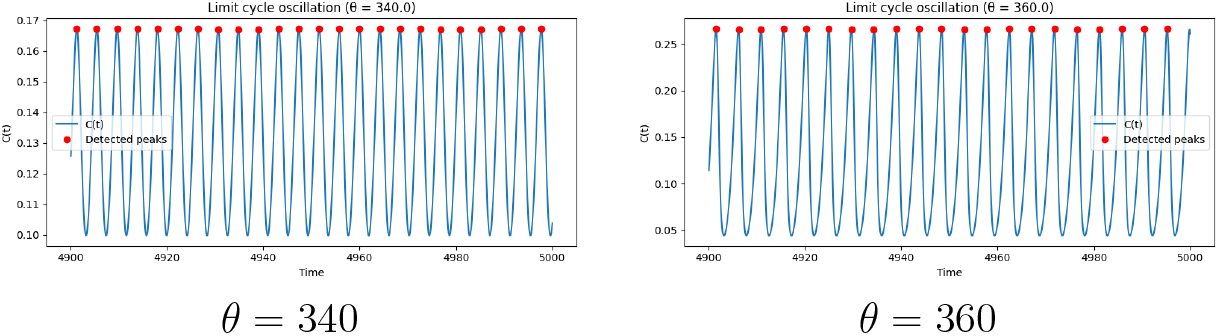
Limit cycle oscillations of tumor cell density *C*(*t*) above the Hopf bifurcation. Each panel shows the last 100 time units of a simulation with total duration *T* = 5000. Red markers indicate peaks used to compute *T*_osc_ and Ω_osc_.

## 4 INCLUSION OF DIFFUSION

To add diffusion back into the above transformation and scaling arguments we recall the linear change of variables from (11):

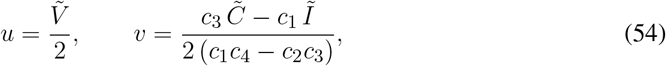

where the coefficients *c*_1_, *c*_2_, *c*_3_, *c*_4_ are given in (17). Thus, *u* depends primarily on the viral perturbation 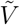, while *v* is a fixed linear combination of the cellular perturbations 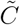 and *Ĩ*.

Since the viral diffusivity was normalized to unity (*D*_*V*_ = 1), *u* inherits diffusion coefficient 1. Similarly, *v* is a linear combination of the two cancer populations, which have a common diffusion coefficient *D*_*c*_ = *D*_*i*_. Hence *v* inherits the same effective diffusivity *D*_*c*_. In the (*u, v*) coordinates the diffusion matrix is diagonal:

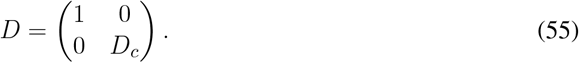

Combining spatial coupling with the linear kinetics, we obtain on the Centre Manifold that

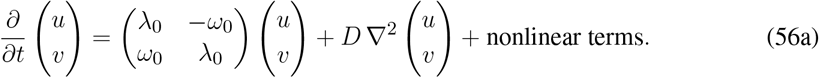

This reduced reaction–diffusion system captures the linear oscillatory modes and their spatial coupling. The next step is to include the weak nonlinearities that saturate these oscillations and ultimately give rise to pattern formation.

### 4.1 Observed Patterns and Explanation

In the previous section we reduced the oncolytic virotherapy model to an oscillatory form, highlighting its proximity to a Hopf bifurcation. Once such a reduction is available, the system can be interpreted within the general theory of coupled oscillators, where diffusion and nonlinear interactions organize local oscillations into coherent spatial structures such as plane waves, spiral waves, and turbulent patterns [19, 12, 38].

In this section, we show numerical simulations of the original *C* − *I* − *V* model (1) using the VisualPDE tool [36]. We explore the dynamics for various values of *θ* and describe the main phenomena that emerge. Throughout these experiments, we impose Neumann boundary conditions. Initial conditions consist of a homogeneous tumor mass (*C, I, V*) = (1, 0, 0) with localized viral inoculation spots, where the concentration is set to (*C, I, V*) = (1, 0, 10). Depending on the experiment, the computational 2-dimensional domain is chosen to be either square or circular with Neumann boundary conditions, as indicated in the figures. These numerical experiments serve as a reference point for the more systematic analysis to follow, where we return to the analytically reduced system to isolate and interpret specific features.

#### Initial Ring Formation and Wave Propagation

To observe the initial behavior upon inoculation, we take circular domain with radius 250 units, and *θ* = 350, with two spots of virus injected. Upon inoculation, the infection spreads in waves of virus chasing the tumor to form expanding circular rings that radiate outward from the source. As shown in Fig. 6, these waves originate at the inoculation site, which for *θ* > *θ*_*H*_, act as a pacemaker sustaining periodic emission of fronts. But over time, we see that these fronts collide, merge, and distort—marking the transition from ordered wave propagation to the irregular patterns to come in later stages.

**Figure 6.**
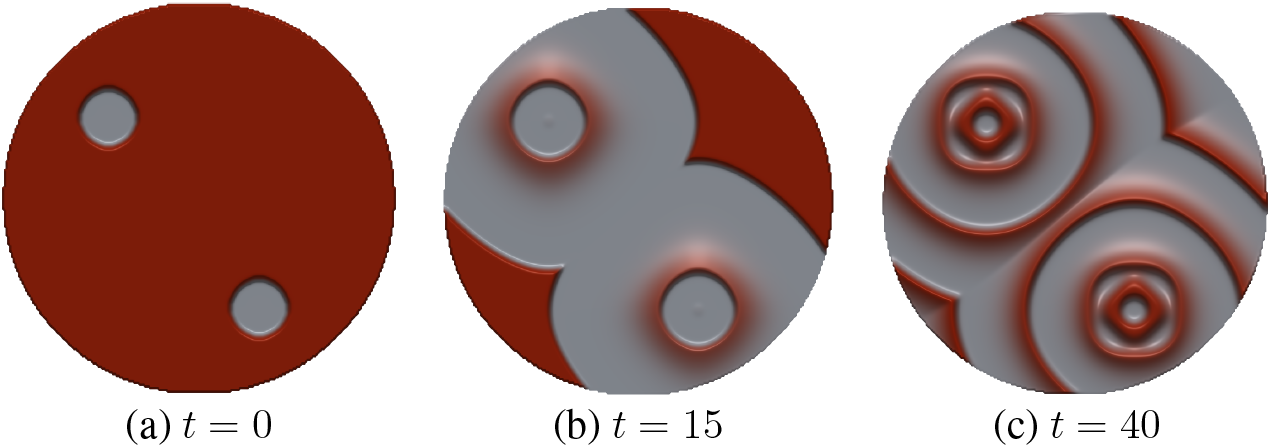
Early dynamics of system (1) following localized viral inoculation (VisualPDE [36]). Domain radius = 250,*θ* = 350, other parameters from Table 1. We have plotted the tumor concentration, where gray regions are close to zero value while the red regions cover high tumor densities. (a) Viral introduction at a point source. (b) Expanding circular fronts form around the core. As fronts widen and interact, curvature decreases and the system progresses toward complex, disordered motion.

#### Spatiotemporal Patterns Beyond Hopf Bifurcation

To see the long term behavior of the system (1), we take a square domain of size 500 units and simulate with viral production rate *θ* = 500, which is well above the Hopf threshold (recall that *θ*_*H*_ ≈ 338). A different color scheme is adopted to make the contrast in tumor density more apparent, where blue region is the lowest, red is intermediate, orange is high and yellow is the highest. Representative snapshots in Fig. 7 show that the domain is tiled by fragmented tumor aggregates whose collisions and mergers sustain long-term turbulent activity rather than relaxing to a simple fixed pattern.

**Figure 7.**
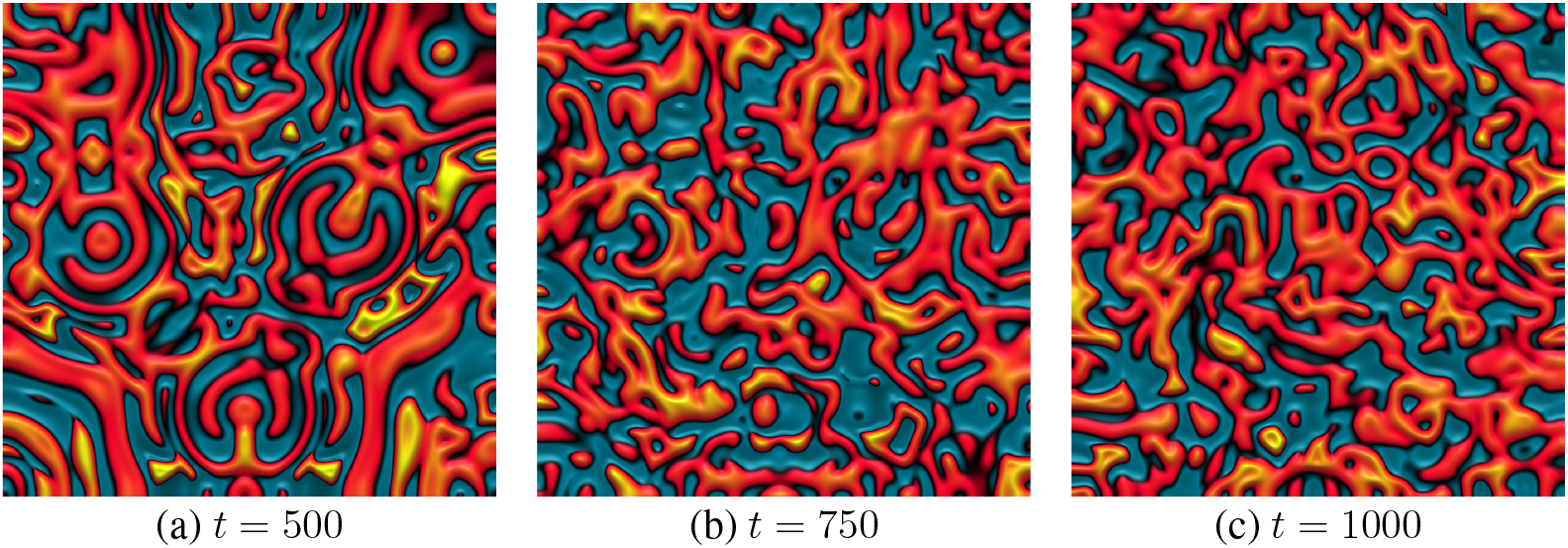
Persistent spatiotemporal activity beyond Hopf bifurcation (*θ* = 500). Snapshots of the field show turbulent organization of tumor mass on a square domain with side 500 units. Simulated using VisualPDE [36].

#### Spiral Fragments

To identify some organizing structure within this turbulent behavior, we consider a close-up sequence in Fig. 8, again for circular domain of radius 250 units and *θ* = 350. In this zoomed in snapshot, we observe a single spiraling disturbance, which seems to rotate and also drift a little, with possibly a length scale for its wavelength and curvature. This serves as a visual clue to look for spiral wave solutions for our original system (1), which we will discuss in more detail in the upcoming sections.

**Figure 8.**
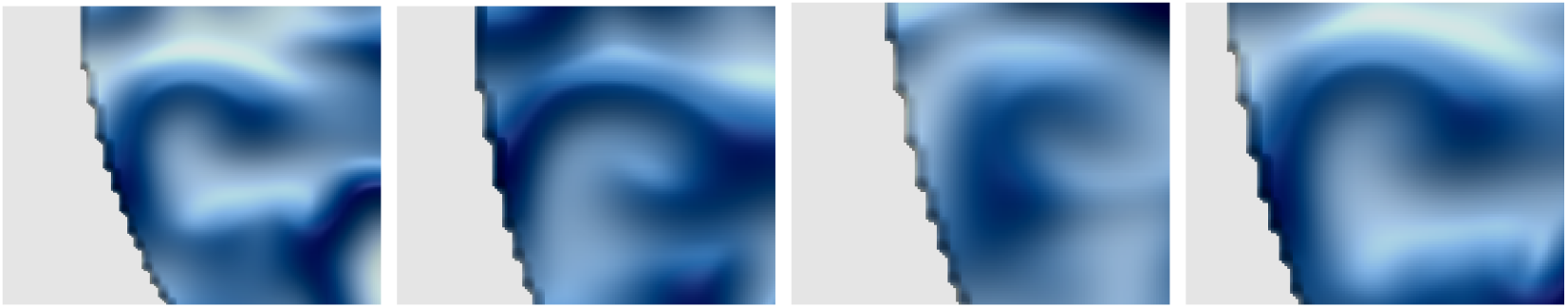
Plot of tumor mass density showing closeup on spiral disturbance for *θ* = 350, with color intensity proportional to tumor concentration. Consecutive frames show a single rotating wave segment where the tip precesses slowly while the arm maintains a rough structure and length scale, characteristic of spiral-wave dynamics (adapted from VisualPDE [36]).

#### Amplitude Modulations at Fixed Spatial Points

To further probe the spatiotemporal behavior, for our choice of parameter *θ* = 350, we restrict attention to the temporal evolution at a fixed spatial point in the domain. Specifically, we record the time series of *V* at *x*_0_ = (10, 20) for a simulation in square domain of side *L* = 100. The resulting dynamics, shown in Fig. 9, reveal a slow envelope riding on top of the fast oscillations near the natural frequency. To better visualize the amplitude modulations, we need to tune out the fast oscillations for which we construct a Poincaré sampling at fixed intervals Δ*t* close to one oscillation period (here, every 27 frames). The resulting trajectory (Fig. 9b) shows roughly how much the amplitude varies over successive return to the same phase of oscillation.

**Figure 9.**
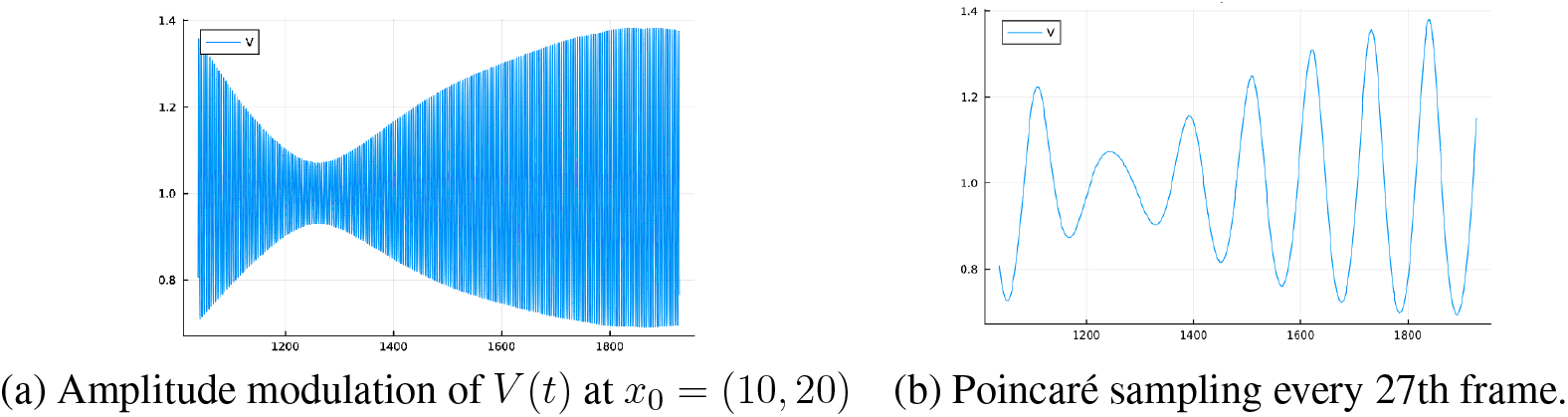
Local temporal structure at a fixed spatial point *x*_0_ = (10, 20) for *θ* = 250 and domain size 100 *×* 100. On the x-axis, we have time and y-axis is *V* = *V* (10, 20, *t*). (a) A slow envelope modulates the fast oscillation near the natural frequency. (b) Sampled once per oscillation, points follow a thin curve clearly showing the amplitude modulations on a time scale of Δ*T* = *T*_*nat*_ ≡ 27 frames.

#### Phase Portraits Across θ

Finally, instead of monitoring the temporal evolution at a single component, we consider the three-dimensional trajectory (*C*(*x*_0_, *t*), *I*(*x*_0_, *t*), *V* (*x*_0_, *t*)) again with fixed *x*_0_ = (10, 20). Sampling again near the natural frequency, which is every 27 frames as done for previous Fig. 9, we obtain the phase portraits shown in Fig. 10. Note that here the axes are normalized so that the unstable fixed point *E*_+_ sits at (1, 1, 1). Closer to the onset of Hopf bifurcation (*θ* = 344), the trajectory forms a relatively thin closed loop in (*C, I, V*); by *θ* = 350, the loop thickens into a torus-like annulus and the instantaneous amplitudes vary more widely.

**Figure 10.**
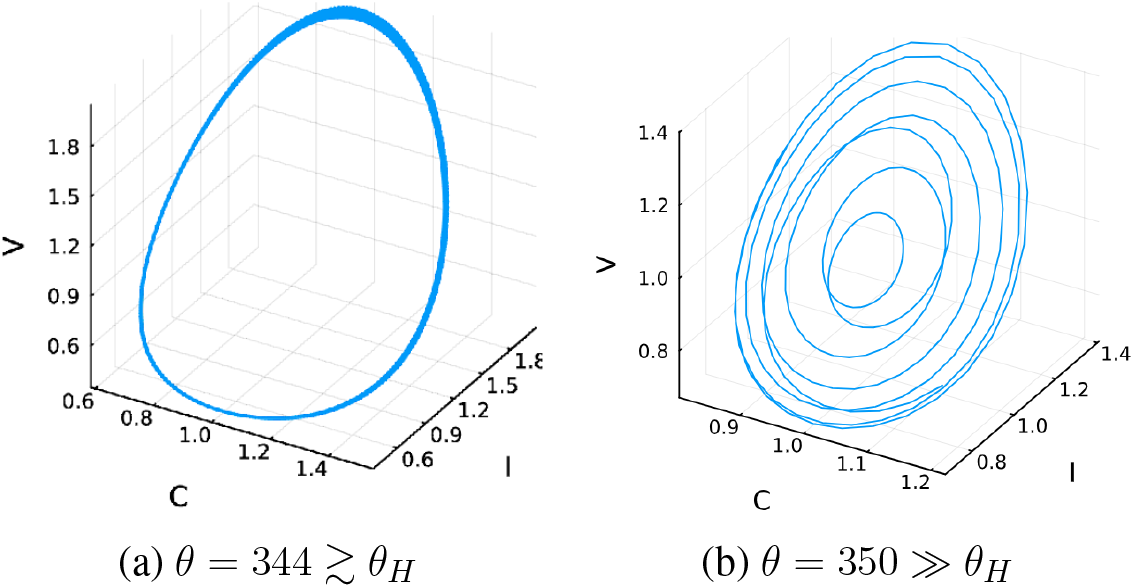
Poincaré phase portraits in (*C, I, V*) sampled near the natural frequency. The *x, y, z* axes are respectively 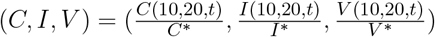, where we normalize the coordinates so that the unstable point *E* _+_ sits at (1, 1, 1). Just beyond onset (*θ*_H_ ≈ 338), trajectories lie on a thin closed loop (approximate limit cycle). At higher value (*θ* = 350), the loop thickens into a torus-like annulus and amplitudes vary substantially.

### 4.2 Explanation

The natural starting point for analysis is therefore to find a family of plane–wave solutions of the cubic complex Ginzburg–Landau equation (CGLE) [2] with real diffusion in two spatial dimensions:

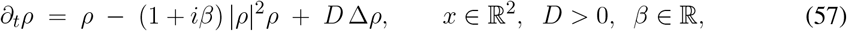

where *ρ*(*x, t*) is the complex amplitude field over the spatial domain and Δ = ∂_*xx*_ + ∂_*yy*_ is the Laplacian.

However, we note that in matrix notation we had *D*_*u*_, *D*_*v*_ for diffusion coefficients but for the amplitude equation, we have a single *D*. Here we take *D* = (*D*_*u*_ + *D*_*v*_)*/*2≈ 0.5 which is supported by the work of Duffy. et al [12], who showed that amplitude equations for plane waves and spiral waves are governed by an effective diffusion rate, which is the average of the two interacting components. But since our goal here is to just explain the patterns and not to make exact predictions, we will be more relaxed with the exact value of *D* such as in spiral wave simulations for next section, where we take *D* = 1.

The complex Ginzburg Landau equation shows a number of spatial patterns, which we consider in sequence.

#### 4.2.1 Spatially uniform oscillations

The spatially uniform oscillation is represented as *ρ*(*x, t*) = *e*^−*iβt*^, which is synchronized oscillation of the whole domain with unit amplitude and frequency *β*.

#### 4.2.2 Plane waves

Take the plane–wave ansatz with amplitude *α* ∈ (0, 1] and wavevector **k** ∈ ℝ^2^,

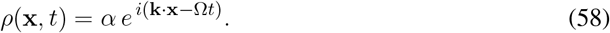

Then |*ρ*|^2^ = *α*^2^, ∂_*t*_*ρ* = −*i*Ω*ρ*, and Δ*ρ* = −|**k**|^2^*ρ*. Substituting into (57) and dividing by *ρ* gives

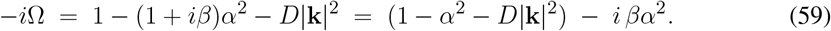

Equating real and imaginary parts, we find

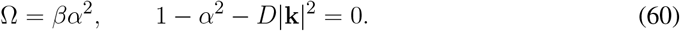

Thus the admissible plane–wave family is

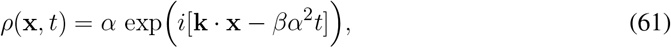

for any wave vector **k** with

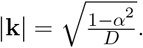

The homogeneous oscillation without spatial propagation corresponds to *α* = 1, for which **k** = 0 and we obtain *ρ*(**x**, *t*) = *e*^−*iβt*^.

Eliminating *α* from (60) gives the **dispersion relation** between frequency Ω and wave mode **k**

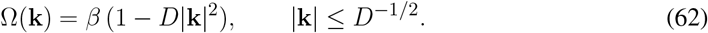

Phase and group velocities are colinear with **k**:

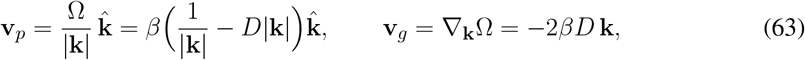

so **v**_*p*_ and **v**_*g*_ always point in opposite directions when **k** ≠ 0.

The plane waves can be seen in the circular spread patterns in Figure 6, where for larger radius, the waves moves in normal direction as a plane wave.

#### 4.2.3 Spiral Waves and Turbulence

Spiral waves present analytic challenges that their plane wave counterparts do not. As noted earlier, spiral dynamics originate at a central core, corresponding to the limit *r* → 0 in polar coordinates centered on the core. This introduces singular terms that must be handled carefully to avoid divergences at the origin, while at the same time ensuring smooth continuity with the far–field plane–wave behavior. Cohen and co–workers [9] developed techniques to bridge this core–to–tail connection, providing the first demonstration that reaction–diffusion dynamics alone can sustain spiral waves without the need for additional terms. Building on this, Duffy et al. [12] formulated a general theory of oscillatory media near Hopf bifurcation, reducing such systems to the *λ*–*ω* class. As Murray’s classic text explains in detail [26], this reduction ultimately leads to the complex Ginzburg–Landau equation (CGLE) (57), which admits both plane–wave and spiral solutions [19].

Comprehensive reviews, most notably the survey by Aranson and Kramer [2], have since documented the full breadth of spiral-wave phenomena, including their stability, the mechanisms of spiral breakup, and transitions to turbulence governed by the cubic CGLE. Returning to Eq. (57), the single control parameter *β* quantifies the strength of amplitude–frequency coupling. The types of spirals that can arise is huge, including linear and logarithmic spirals as well as one-armed, two-armed or *m*-armed spirals, and meandering sprials. All these are well documented in the cited literature [9, 12, 26, 2, 33] and we will not recall the underlying scaling arguments here. Instead, we reproduce the main spiral regimes through direct numerical simulations and compare them with the theoretical expectations.

### 4.3 Numerical simulation of spiral waves

To illustrate the spiralling solutions of Eq. (57), we perform direct numerical simulations on a two–dimensional square domain, with Neumann boundary conditions. The domain size is *L* = 100 with *N* = 256 grid points in each direction, giving a spatial resolution Δ*x* = *L/N* ≈ 0.39. The Laplacian is discretized using second–order central differences, and time integration is carried out with an explicit Euler scheme with timestep Δ*t* = 10^−3^. To visualize the spiral waves we have chosen an explicit solver instead of VisualPDE, since an explicit solver allows us full control on the discretization, time stepping, and initial condition. Our choices above are sufficient to resolve the spiral dynamics without visible numerical artifacts.

To seed a spiral, we prescribe

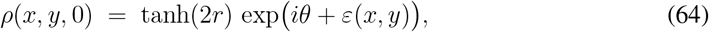

where 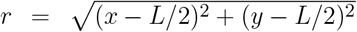 is the distance from the domain center and *θ* = arctan 2(*y* − *L/*2, *x* − *L/*2) the polar angle. The prefactor tanh(2*r*) creates a smooth core, while the phase factor *e*^*iθ*^ winds the oscillation once around the center, imposing a single spiral seed. A small random perturbation *ε*(*x, y*) ∼ 𝒰 (− 10^−3^, 10^−3^) is added to break perfect rotational symmetry and prevent numerical artifacts.

To interpret the spiral dynamics of (57), we monitor both the amplitude |*ρ*(*x, y, t*)| and the phase arg(*ρ*(*x, y, t*)). In practice, we display the amplitude defect 1 − |*ρ*|^2^ as a color map, which highlights spiral cores as bright spots, and overlay contours of constant phase to reveal the rotating arms and their interactions. This joint representation makes it possible to simultaneously follow the wavefront geometry and the location of topological defects. An example is shown in Fig. 11.

**Figure 11.**
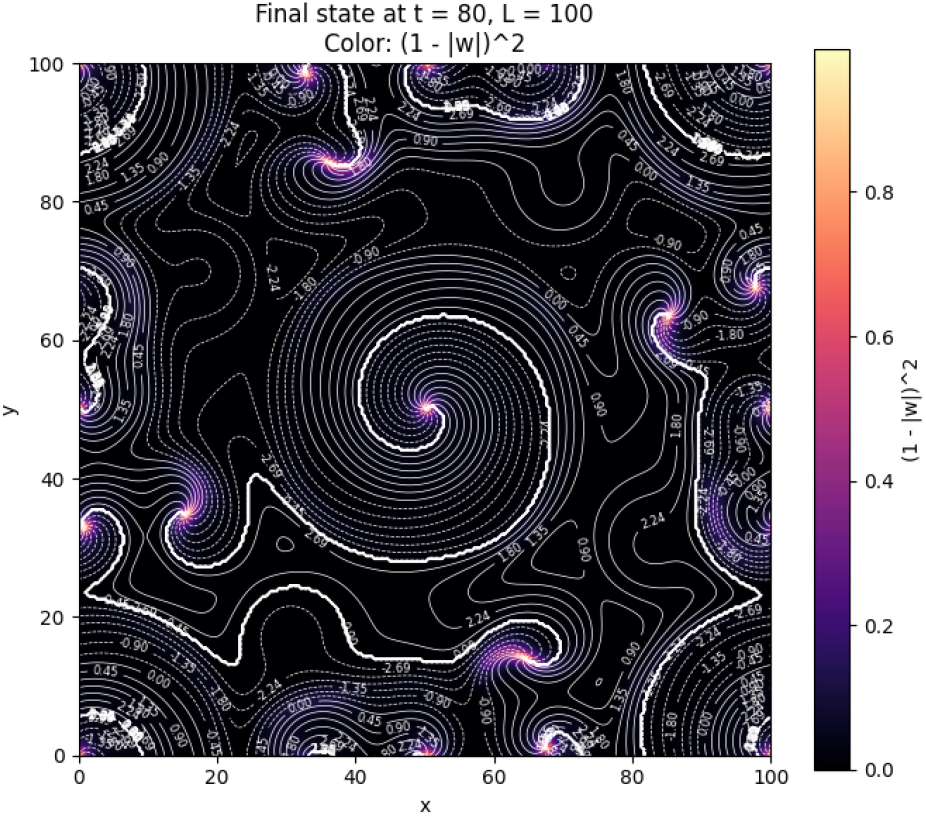
Spiral wave dynamics for CGLE (57) for *β* = 1.5 and *D* = 1 at *t* = 80*s* in square domain of size 100 obtained from the seeded initial condition (64). Color shading shows the amplitude defect (1− |*ρ* |^2^), making spiral cores visible as bright spots, while overlaid contours trace equal phase lines of the oscillation, revealing the structure of the rotating arms. The initial seed had one spiral at the centre, but many more spiral centres arise over time.

### 4.4 Onset of turbulence

The dynamics of Eq. (57) depend strongly on the amplitude–frequency coupling parameter *β*. For small values of |*β*|, the medium supports *frozen spirals*: coherent, steadily rotating structures with smooth arms (Fig. 12(a)). Far from the spiral core, the wavefronts match the plane–wave dispersion relation (62) derived in the previous section, while the core itself serves as a continuous source of rotational disturbances.

**Figure 12.**
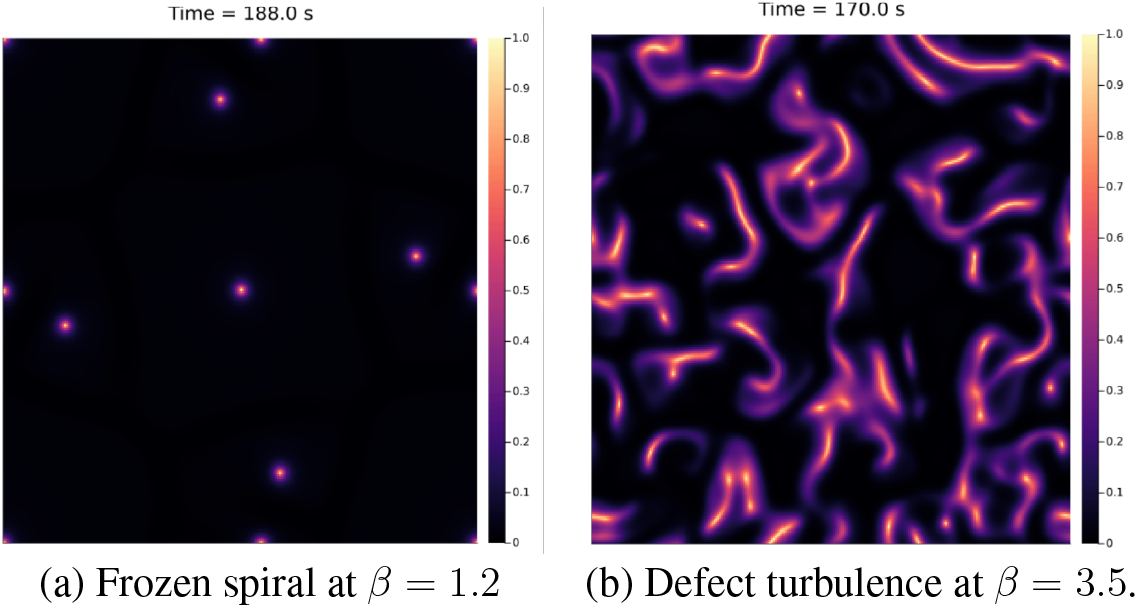
Spiral wave states in the CGLE (57) with *D* = 1. (a) At small *β*, the system sustains frozen spirals: stable rotating waves with coherent arms. (b) At larger *β*, nonlinear frequency shifts destabilize the arms, leading to spiral breakup and a turbulent regime with proliferating defects.

As |*β*| increases, nonlinear frequency shifts destabilize the spiral arms and it becomes harder to sustain synchronous oscillations across domains of differing amplitude. The wavefronts begin to develop irregularities, tips break apart, and topological defects proliferate, ultimately leading to *defect turbulence* (Fig. 12(b)).

Here we refer to a more recent work by Szczesny et al. [33], which studied spiral-wave stability in spatial rock–paper–scissors models (which also reduce to the same CGLE we consider here). They identified three regimes: (i) the *bound state (BS)* phase of long-lived frozen spirals (small *β*, Fig. 12(a)); (ii) the *Eckhaus instability (EI)* phase, where far-field perturbations destabilize spiral arms but the core remains intact; and (iii) the *absolute instability (AI)* phase, where instability propagates into the core and prevents any sustained spirals. Our simulations (Fig. 12) reproduce this sequence—from frozen spirals at low *β* to turbulence at larger *β*. However, we cannot distinguish between the EI and AI phases in our numerical experiments.

## 5 CONCLUSION

To summarize, we started with a basic Reaction-Diffusion (RD) model for the replication and spread of an oncolytic virus in a tumor population, treating the viral burst size as the primary control parameter. We specifically wished to explore and explain the patterns that arise right above the Hopf bifurcation threshold, which we achieved through systematic reduction to the Complex Ginzburg–Landau Equation (CGLE) (57). In light of the normal form approach taken here, we identified the parameter *β*, which measures the asynchrony of oscillations (the degree to which frequency depends on amplitude), as the critical factor. It is known through previous work on amplitude-dependent frequency oscillators, and we also saw through our simulations, that this *β* term is responsible for the stability of different patterns in the medium.

Similar behavior in oncolytic virotherapy has been observed before in much more complicated models involving immune interactions [31], evolving resistance [6], agent-based simulations [40], detailed tumor geometries [7], and nuanced movement strategies (like pressure-driven flow) [25]. However, we place the importance of our work in its simplicity: for a treatment that is inherently oscillating, the implications on the spatiotemporal patterns can be identified by *β*, a parameter that is both simple to describe and easy to calculate. Specifically, if the eigenvalue at the unstable point is *λ*_0_ + *iω*_0_ and the frequency of the resulting limit cycle is *ω*, then *β* is proportional to the difference in frequencies relative to the distance from the threshold. The parameter *β* is proportional to 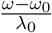. It is known that if *β* is smaller than a certain critical value (around 3, depending on the system), we can expect stable spiral patterns, while higher values of *β* lead to turbulent patterns [2] as the general outcome. However, Wodarz et al. [40] found a very curious result in their experiments, where the same system with the same initial conditions could lead to different treatment outcomes. They attributed this unpredictability either to intrinsic stochasticity within the model or to external anti-viral factors not included in the model [40], and the same problem persists in our model too, but it is also the natural side effect of reducing the dynamics too much.

Finally we reconsider the mathematical approach taken here, which reduced the complex dynamics to the picture of coupled nonlinear oscillators. This could be used to explain the main patterns already, but we can go further. In particular, a previous study [3] found that an inoculated spot of virus can split into two at regular intervals, a phenomenon termed periodic spot-splitting. We can now offer a possible explanation for this using the idea of amplitude-dependent frequency of the oscillators. In the initial inoculation, the center of the spot has the highest viral load (amplitude), while a point near the edge has a lower value. If the oscillators were truly linear, we would expect the entire spot to recover simultaneously after one oscillation. However, because we have shown the system is governed by the nonlinear Stuart-Landau Equation, the frequency of the highly infected spot center will be different from the less infected off-center points. This leads to a growing phase gap between the center and the periphery. This phase difference could create a scenario where the center of the spot reaches its minimum biomass (near-extinction) while some off-center point is still near its maximum—a desynchronization that would manifest visually as the splitting of the original inoculated spot. While this explanation remains heuristic, it clearly motivates a direction for further mathematical inquiry: establishing the precise conditions on the parameter *β* that determine the onset, choice of direction for symmetry breaking and the subsequent dynamics of spot splitting, thereby contributing to understanding of this new type of pattern formation for its own sake.

To conclude, we provide here a robust, minimal mathematical framework for understanding the failure modes of oncolytic virotherapy in spatially extended tumors. Our analysis demonstrates that, while constant efforts are made to increase viral contagiousness, high replication rates are mathematically destined to lead to fragmented patterns. This inherent instability ensures unreliable and incomplete viral coverage, confirming that maximizing viral replication still leads to therapeutic failure [27]. This finding supports the existing consensus that oncolytic viruses due to their dynamic limitations, cannot reliably clear a tumor entirely on their own and should therefore be strategically used in combination with other therapies [37, 15, 24]. This necessitates a shift from monotherapy optimization to designing viruses from scratch for synergistic action within combination treatment strategies.

## CONFLICT OF INTEREST STATEMENT

The authors declare that the research was conducted in the absence of any commercial or financial relationships that could be construed as a potential conflict of interest.

## AUTHOR CONTRIBUTIONS

TB performed the analysis, the numerical simulations, and wrote the first draft. TH conceptualized the study, oversaw its development, and contributed to the final write-up.

## FUNDING

TH is supported through a Discovery Grant of the Natural Science and Engineering Research Council of Canada (NSERC), RGPIN-2023-04269.

## ACKNOWLEDGMENTS

TB and TH are grateful to helpful comments from A. Baabdulla, N. Mohammadnejad, M. van Walsum, and the Mathematical Biology Journal Club. TB is also thankful for 2025 Gene Golub SIAM Summer School, and its organizers Ryan Goh and Jason Bramburger for all their help.

## Notes

### Competing Interest Statement

The authors have declared no competing interest.

